# Seipin Regulates Caveolin-1 Trafficking and Organelle Crosstalk

**DOI:** 10.1101/2024.09.17.613438

**Authors:** Maxime Carpentier, Mohyeddine Omrane, Rola Shaaban, Jennica Träger, Naima El Khallouki, Mehdi Zouiouich, Marie Palard, Takeshi Harayama, Corinne Vigouroux, Soazig Le Lay, Francesca Giordano, Xavier Prieur, Abdou Rachid Thiam

## Abstract

Caveolin-1 (CAV1), the main structural component of caveolae, is essential in various biological processes, including mechanotransduction, lipid metabolism, and endocytosis^1–4^. Deregulation of CAV1 dynamics is linked to various pathologies, including cellular senescence, cancer, insulin resistance, and lipodystrophy^5–9^. However, mechanisms regulating CAV1 trafficking and function remain poorly understood. Here, we show that seipin, a crucial lipid droplet (LD) biogenesis factor^10^, modulates CAV1 trafficking. Deletion of seipin resulted in the accumulation of saturated lipids, leading to ceramide and sphingomyelin overproduction, which disrupted the membrane order of the trans-Golgi network (TGN). In seipin deficiency, CAV1 location to the plasma membrane (PM) was impaired, reducing caveolae. Instead, CAV1 accumulated in TGN and late endosome compartments, which fused with LDs and delivered the protein. In wild-type (WT) cells, this process was minimal but significantly enhanced by treatment with palmitate, ceramide, or Stearoyl-CoA desaturase-1 (SCD1) inhibition. Conversely, in seipin-deficient cells, inhibiting Fatty Acid Synthase (FASN) or overexpressing SCD1 restored CAV1 localization to the PM and reduced its accumulation in LDs. Our findings reveal that seipin controls the funneling of palmitate toward glycerolipids synthesis and storage in LDs versus conversion to ceramides in the ER. This balance is crucial to cellular protein trafficking by controlling the TGN membrane order. Therefore, our study identifies seipin as a critical regulator of cellular lipid metabolism, protein trafficking, and organelle homeostasis. These findings shed light on the processes regulating CAV1 trafficking and show that convergent pathophysiological mechanisms associated with defects in CAV1 and seipin contribute to metabolic disorders, including insulin resistance and lipodystrophies^11–14^.

## Introduction

Cells respond to metabolic demands by adjusting their lipid metabolism, which is essential for maintaining proper cellular function. This regulation involves lipids’ uptake, secretion, and storage, leading to dynamical changes in cellular membrane lipid composition and mechanical properties to meet metabolic needs^15,16^. Dysregulation in these processes is implicated in various disorders, including Type 2 diabetes, obesity, lipodystrophy, and cancers. One of the major players in these processes is Caveolin-1 (CAV1)^5,17,18^, linking membrane mechanics to metabolism^15^. CAV1’s mechanosensing properties through caveola formation are critical for buffering PM surface tension fluctuations and influencing diverse signaling pathways^19,20^. Additionally, CAV1 regulates the transport of lipids and enzymes from caveolae to different organelles^21–23^, which are essential for the function of the organelles. However, the mechanisms by which CAV1 traffics or transports these to their destination organelle remain elusive.

CAV1 regulates the translocation and levels of the Glucose Transporter Type 4 (GLUT4)^14,24^, the Cluster of Differentiation 36 (CD36)^22,25^, and other transporters^26,27^, indirectly or directly, by incorporating them into caveolae. This regulation impacts energy intake, storage, and utilization. Null biallelic variants in *CAV1* are responsible for congenital generalized lipodystrophy type 3 (CGL3)^12^, characterized by a near-total absence of adipose tissue, lipid storage defects, insulin resistance with metabolic complications, and esophageal neuropathy^8,12^. Similarly, loss-of-function mutations in *BSCL2,* encoding seipin, which regulates LD formation and function, result in congenital generalized lipodystrophy type 2 (CGL2), characterized by similar insulin resistance-related metabolic complications associated with cognitive defects^13^. Yet, despite the roles of CAV1 and seipin in lipid metabolism, whether there is a link between the two proteins is unknown. Given that the dysfunction of either protein leads to similar metabolic signatures and neurological involvement, we hypothesized the existence of a functional axis linking seipin and CAV1. We investigated such a potential link since it could provide critical insights into lipid metabolism.

## Results

### Seipin deletion perturbs CAV1 dynamics and caveola formation

As the glucose and fatty acid transporters’ function relies on CAV1, we asked whether seipin removal impacts these enzymes in HeLa cells, which respond to insulin. We cultured WT and seipin KO cells (SKO) in the presence of insulin and transfected them with GLUT4-mCherry. In WT cells, GLUT4 localized to the PM, and in GLUT4 vesicles having a perinuclear localization (Fig. 1A-B). In contrast, GLUT4 PM localization was much lower in the SKO cells, and GLUT4 vesicles were dispersed throughout the cell. In response to insulin, GLUT4 vesicles fuse with PM using SNARE components, including SNAP23 and VAMP2. Upon insulin stimulation, both SNAREs (Fig. S1A-B and S1D-E), especially SNAP23, had reduced recruitment to the PM in SKO compared to WT, agreeing with the defect in GLUT4 PM translocation. This finding directly links seipin deficiency and insulin resistance. CD36 PM localization was also altered in SKO cells (Fig.S1C and S1F; See Material and Method for PM signal quantification).

**Figure 1:**
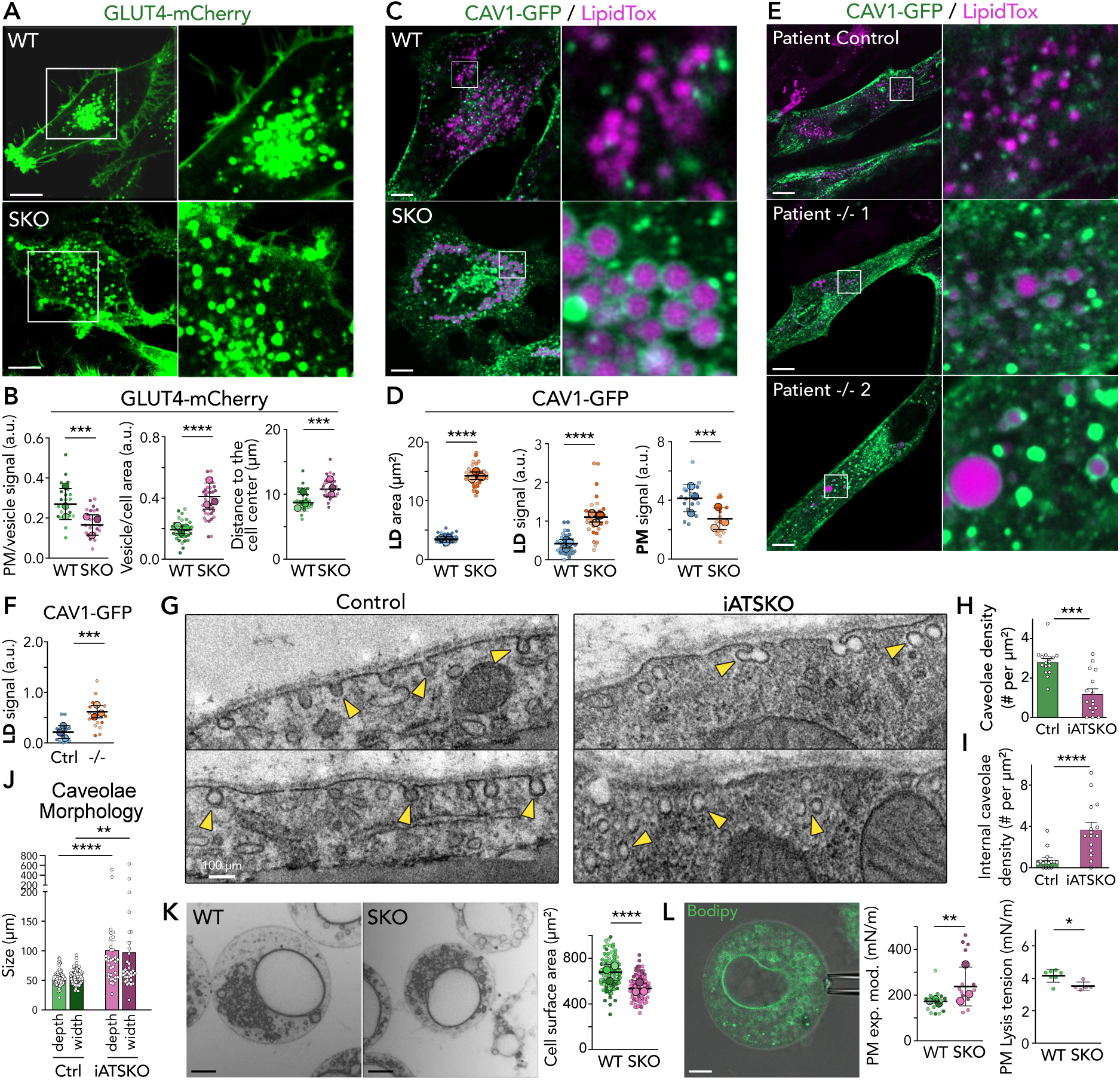
Seipin depletion perturbs CAV1 dynamics and caveola formation. **A.** Representative images of HeLa WT and SKO cells overexpressing GLUT4-mCherry. **B.** Quantification of A. Left: PM signal normalized by the average signal of GLUT4 vesicles; P = 0.0007, n = 21 for WT, 22 for KO; 3 replicates. Middle: sum of vesicle area normalized by cell area; P < 0.0001, n = 35 for WT, 34 for KO; 3 replicates. Right: GLUT4 vesicles’ dispersion in the cell; two-tailed Mann-Whitney analysis, P = 0.0011, n = 34 for WT and 24 for KO; 3 replicates. **C.** HeLa WT and SKO cells overexpressing CAV1-GFP. LDs are induced after transfection by oleate supply. LipidTox Deep Red stained LDs. **D.** Left and middle: mean LD area and normalized CAV1-GFP signal around LDs per cell in C. A two-tailed Mann-Whitney analysis is done with P < 0.0001 for both. For the LDs’ area, n = 61 for WT cells and 39 for KO cells, 3 replicates. For CAV1-GFP signal around LDs’ quantification, n = 30 for WT and 29 for KO cells; 3 experiments. Right: mean CAV1-GFP PM signal measured from swollen cells normalized by basal cell signal; P = 0.0010, n = 15 for WT and KO cells; 3 replicates. **E.** Fibroblasts from control and CGL2 patients overexpressing CAV1-GFP. LDs are induced after transfection by oleic acid supply. **F.** Normalized CAV1-GFP signal around LDs per cell for experiments in E. P = 0.0009, n = 15 cells for each cell line; 3 experiments. **G.** Transmission electron microscopy of cells from control and iATSKO mice. Yellow arrowheads depict caveola vesicles. Scale bar: 100 nm. **H, I, J.** Quantification of Mean Caveolae density per cell, mean internalized caveolae density, and width and depth, respectively. For caveolae density quantifications, a two-tailed Mann-Whitney analysis is done with respective P-values of 0.0002 and < 0.0001; n = 16 for CTLR and iATSKO. For caveolae morphology, a two-tailed Mann-Whitney analysis is done with P-values < 0.0001, for n = 16. **K.** Left: HeLa WT and SKO cells are submitted to a hypotonic medium with Bodipy 493/503 added. A negative color mode is used. Right: quantification of the swollen WT and SKO area from a median plan image. P < 0.0001, n = 120 for WT and KO cells; 3 experiments. **L.** Left: Example of a swollen HeLa cell manipulated with a micropipette. Right: Quantification of the apparent area expansion modulus and lysis tension of swollen WT and SKO cells. Two-tailed Mann-Whitney analysis, top (P = 0.061 and 0.0303; n = 13 for WT and KO cells), bottom (n = 7 WT cells and 5 KO cells); 3 experiments. Unless indicated, scale bars are 5 µm.

CAV1 is one of the major proteins involved in lipid intake, especially in intestinal cells for dietary fat adsorption or adipocytes for lipid storage^28^. Dysfunction of CAV1 perturbs the dynamics of GLUT4 and CD36 vesicles, amongst other transporters localizing at the PM^21,25,26^. CD36 partitions presumably to CAV1 caveolae^22,25^. Thus, we asked if CAV1 localization was impacted by seipin expression in HeLa cells.

In WT cells, overexpressed CAV1 localized mainly to the PM, but this localization was significantly diminished in SKO cells (Fig. S1G). Since CAV1 interacts with cholesterol^29^, we expressed the D4H cholesterol probe and found it much less recruited to the PM in SKO (Fig. S1H-I).

In WT cells loaded with oleic acid to induce LD formation, endogenous and overexpressed CAV1 correctly targeted the PM (Fig.1C-D, S1J-M). Additionally, CAV1 displayed signals in cytosolic vesicles and was rarely localized to LDs, contrasting with Caveolin 2 and 3 (Fig. S1N-Q). In SKO cells, CAV1 failed to target the PM but was surprisingly relocalized to LDs (Fig.1C-D). Compared to CAV1, over-expressed Cavin1 (Fig. S1R), the other key player in caveolae formation, did not accumulate around the LDs. The PM localization of CAV1 in SKO cells was rescued by overexpressing seipin-mScarlet, which also reduced the localization of CAV1 to LDs (Fig.S1S-U). These findings were corroborated in skin fibroblasts from healthy and patient donors with seipin deficiency. Specifically, in the cells from patients, we found less CAV1 PM signal and strong LD binding phenotypes (Fig.1E-F, S2D). These data suggest that seipin regulates CAV1 trafficking.

To further study this impact in the context of adipocytes, we analyzed the adipose tissue (AT) of mice carrying an inducible seipin deletion in adipocytes upon tamoxifen addition (iATSKO). Electron microscopy (EM) analyses revealed a marked decrease in the number and density of caveolae in iATSKO adipocytes, accompanied by increased internal vesicles (Fig. 1G-J, S2A). This result echoed the cell data, showing a decrease in CAV1 at the PM and its accumulation in intracellular vesicles. Thus, our data further support the role of seipin in regulating CAV1 localization and caveola formation.

Finally, considering the role of CAV1 in buffering PM surface tension to withstand mechanical stresses, we subjected both WT and SKO cells to the same hypotonic medium for 10 min. This treatment caused cell swelling and PM stretching. We observed that WT cells were, on average, significantly larger than SKO cells, likely due to the loss of caveola membrane reservoirs upon seipin deletion (Fig.1K, S2B). Additionally, the PM of WT cells exhibited a lower apparent area expansion modulus and a higher lysis tension (Fig. 1L, S2C). These results indicate that seipin dysfunction impairs caveola and PM membrane mechanics.

### Seipin depletion perturbs the lipid composition of the TGN

CAV1 exits the ER in assembled 8S structures, which then relocate to the Golgi apparatus. At the TGN, these structures assemble into higher-oligomerized 70S ones, subsequently transported to the PM^30,31^. A dysfunction in one of these steps could explain CAV1 mistargeting the PM. We used the retention using selective hooks (RUSH) assay^32^ with a GPI-anchored protein to investigate if the secretory pathway was impaired. Trafficking of the GPI protein from the ER to the Golgi was initially similar between WT and SKO cells following biotin addition (Fig. 2A,2B). After 1 hour, the GPI protein was almost undetectable in the Golgi in WT cells and had reached the PM, whereas it remained stuck in the Golgi in SKO cells (Fig. 2A,2B). This data argues that protein movement from the TGN to the PM was altered in SKO cells.

**Figure 2:**
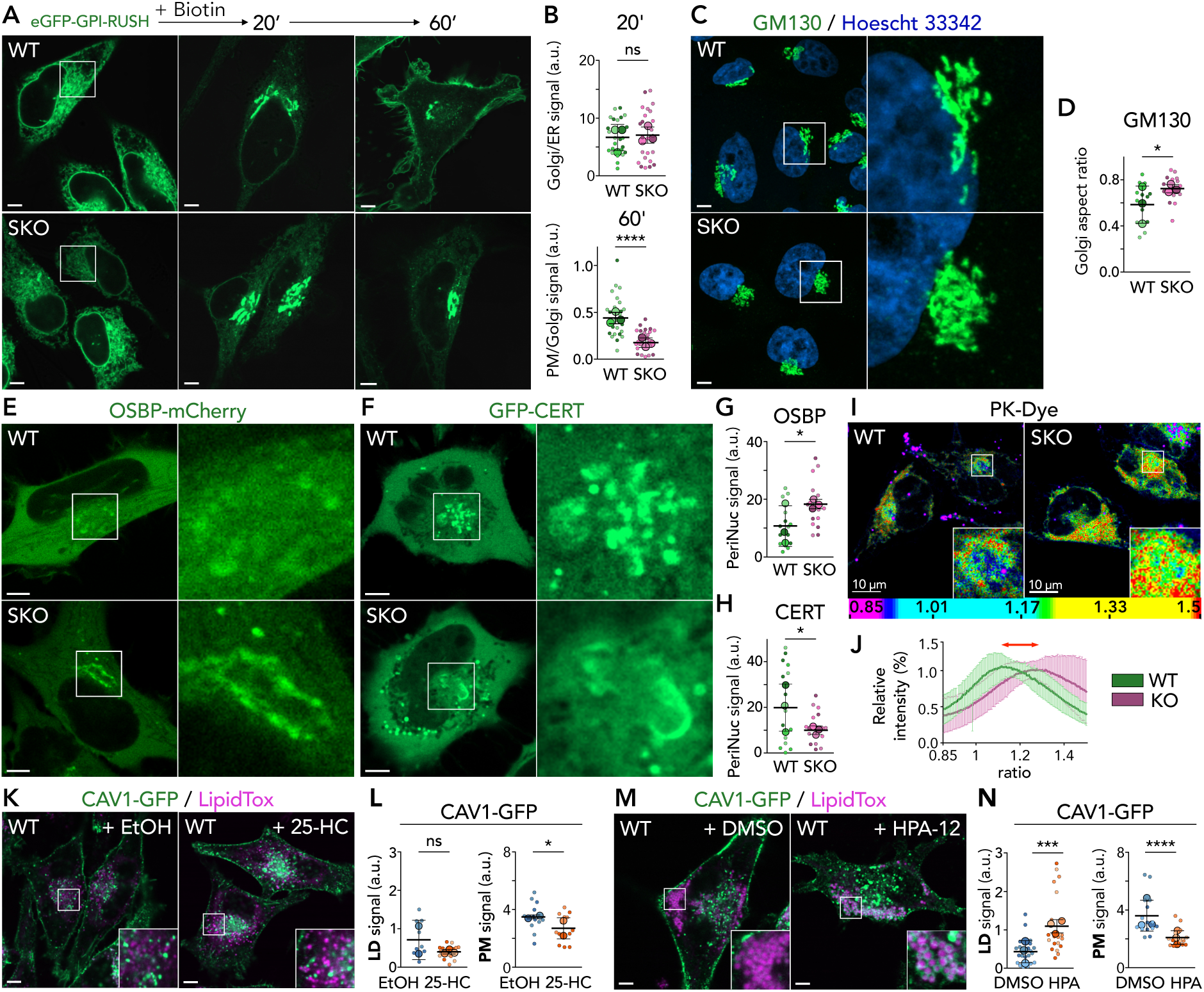
Seipin depletion perturbs the lipid composition and function of the TGN. **A.** HeLa WT and SKO cells overexpressing an eGFP-GPI-RUSH are imaged before and 20 and 60 minutes after biotin addition. **B.** Top: the mean signal ratio between the Golgi and the ER at 20 minutes. P = 0.7278; 3 experiments (8 cells for each). Bottom: the mean signal ratio between the PM and the Golgi at 60 minutes. P < 0.0001, n = 24 cells for each; 3 replicates. **C.** Immunofluorescence of WT and SKO HeLa cells against Golgi resident protein GM130 and labeled with Hoechst. **D.** Golgi aspect ratio measuring the ratio between the largest Golgi particle’s minimum and maximum Feret diameter. P = 0.0136, n = 15 each; 3 experiments. **E, F.** WT and SKO HeLa cells overexpressing the cholesterol transporter OSBP-mCherry or the ceramide transporter GFP-CERT. **G**, **H.** A mask of the Golgi apparatus was created from an average Z-project, and the sum of the integrated intensity from the Golgi particle mask was measured and divided by the mean cytoplasmic signal. Panel G: P = 0.0127, n = 15 cells for each condition; 3 experiments. Panel H: P = 0.0472, n = 15 cells for each case; 3 replicates. **I.** Ratiometric images of PK-Dye incubated with HeLa WT and SKO cells. **J.** Relative intensity (%) associated with the corresponding ratio value from experiments in I; n = 24 for WT and 37 for KO cells; 3 experiments. **K.** HeLa WT overexpressing CAV1-GFP and oleate-loaded. A supply of 2.5 µM of 25-Hydroxycholesterol (25-HC) is done during the transfection and maintained afterward. Loading control is done with Ethanol (EtOH). **L.** Left: mean normalized CAV1-GFP signal around LDs per cell. Two-tailed Mann-Whitney analysis is done, P = 0.1303, n = 10 cells for WT + EtOH from 2 replicates and 15 cells for WT + 25-HC from 3 replicates. Right: mean CAV1-GFP PM signal. P = 0.0463, n = 12 cells for WT EtOH and 14 cells for WT + 25-HC from 3 replicates. **M.** HeLa WT cells overexpressing CAV1-GFP and with LDs induced. A loading of 2.5 µM of HPA-12 is done during the transfection and maintained afterward. Control is done with DMSO. **N.** Left: mean normalized CAV1-GFP signal around LDs per cell. Two-tailed Mann-Whitney analysis, P = 0.0001, n = 21 cells for each condition from 3 replicates. Right: mean CAV1-GFP PM signal. Two-tailed Mann-Whitney analysis, P < 0.0010, n = 15 cells each from 3 experiments. Scale bars: 5 µm.

We stained GM130 to check the integrity of the Golgi and found that it was rounder for SKO cells (Fig. 2C-D). The Golgi morphology was also altered in fibroblast cells from patients with CGL2 compared to healthy donors upon oleic acid supply (Fig. S3A-B). The TGN has a lateral organization critical for cargo sorting and trafficking to the PM and other destinations^33,34^. This organization includes forming lipid-ordered domains rich in cholesterol and sphingomyelin (SM). Cholesterol is brought by the Oxysterol-binding protein (OSBP) from the ER and counter-exchanged by the retrieval of PI4P^35^ at TGN-ER contact sites. SM is synthesized by sphingomyelin synthase, which converts ceramides transported from ER^36^ by the ceramide transfer protein (CERT). These structured TGN regions are presumably crucial for sorting the 70S structures, which are rich in cholesterol^29^.

In SKO cells, overexpressed OSBP was stuck at the TGN-ER interface compared to WT cells, indicative of a defect in its transport activity and the accumulation of PI4P in the TGN^37^ (Fig. 2E,2G). Accordingly, the PI4P P4M reporter was more intensely present in the perinuclear area^7^ (Fig. S3C, S3E). Additionally, overexpressed CERT (Fig. 2F-H), which had a focused perinuclear localization in WT cells, displayed a scattered localization in SKO cells. Lysenin, an SM marker, also showed a scattered SM signal, which was less concentrated in the Golgi than in WT cells (Fig. S3D, S3F).

These data suggest an impaired TGN architecture and cholesterol and SM composition. Therefore, we used the Prodan-based PK dye membrane order reporter to analyze membrane order. We found a decrease in order in the perinuclear region, supporting that the TGN membrane order was reduced (Fig. 2J-K). Lipidomics of Golgi-enriched fractions showed on average a decrease in SM and cholesterol levels (Fig. S3G-H).

The alteration of the TGN order in SKO cells could explain the mislocalization of CAV1 to LDs at the expense of the PM. Thus, perturbing TGN membrane order in WT cells could replicate the SKO phenotype. To test this, we inhibited OSBP or CERT transfer activity in WT cells by supplying 25HC or HPA12, respectively. 25HC supply resulted in a slight decrease in the PM signal of CAV1 (Fig. 2K-L, S3I). CAV1 vesicles were more frequently recruited near LDs, but the protein signal was on the LD surface. In contrast, CERT inhibition phenocopied SKO cells as the CAV1 signal at the PM decreased, and the protein was found on the LD surface (Fig. 2M-N, S3J). These data support the idea that the TGN membrane lipid order regulates CAV1 trafficking to the PM or LD.

### CAV1 relocalizes from the TGN/Late endosomes to LDs

In WT cells, unlike CAV2 and CAV3, newly translated CAV1 did not target LDs from the ER, probably because it rapidly exits it to prevent aggregation of its 8S complex^30^. In the SKO cells, CAV1 was absent from the ER and was on vesicles docked to LDs (Fig. S4A). This evidence argues that CAV1 did not relocalize to LDs from the ER.

The trafficking defect of CAV1 between the TGN and PM was concomitant with the accumulation of CAV1 vesicles in the cytoplasm, often close to LDs. We hypothesized that these vesicles contained TGN material carried to maturing endosomes. RAB5, an early endosome marker, rarely colocalized with CAV1 and LDs in WT or SKO cells (Fig. S4B). On the other hand, RAB7 vesicles, marking late endosomes, were more frequently enriched in CAV1 and in proximity to LDs in SKO than WT cells (Fig. 3A-B, S4C). Agreeing with our hypothesis, lysosomal components were also recruited to LDs in SKO cells (Fig. 3C, S4D-E).

**Figure 3:**
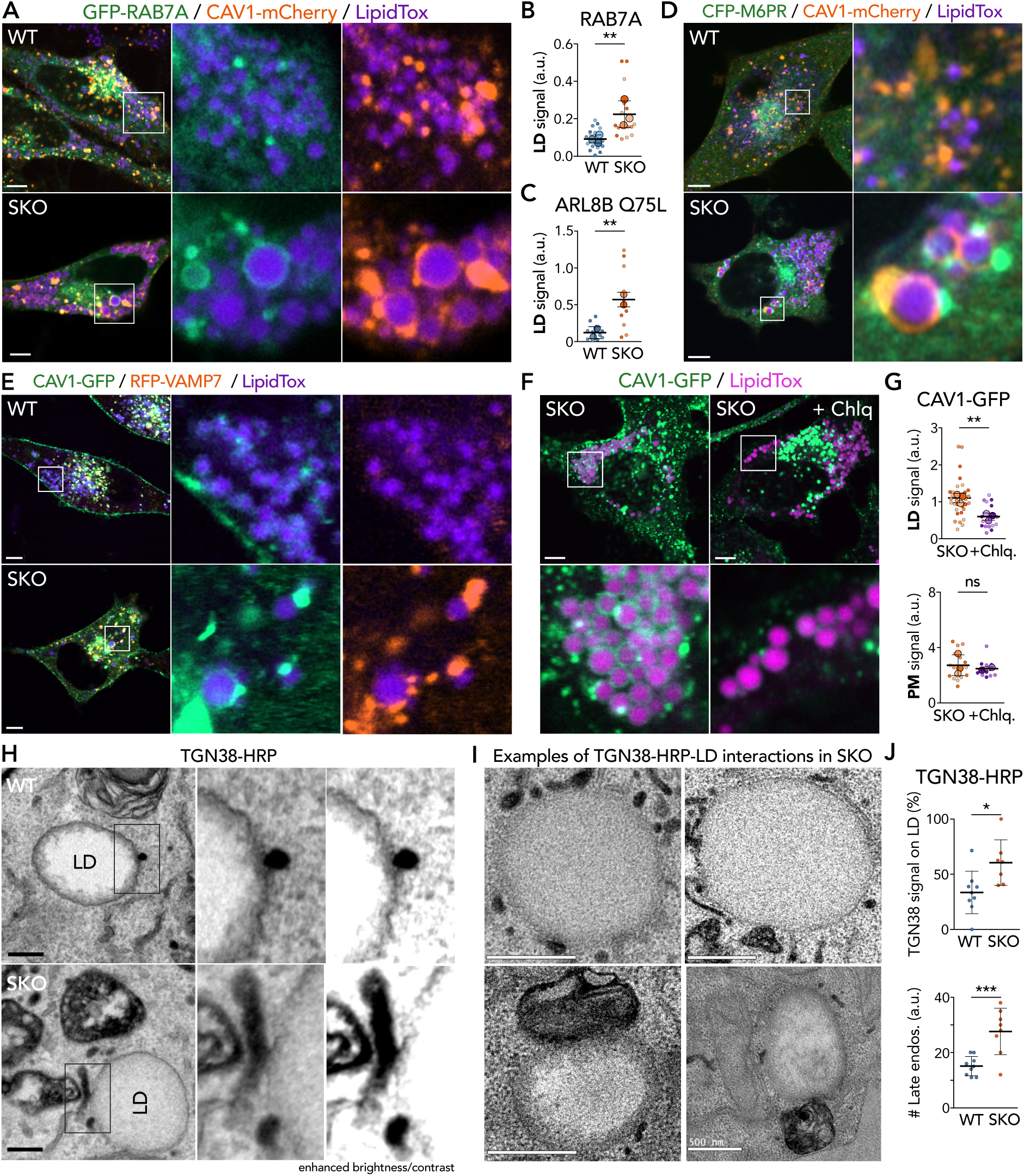
CAV1 relocalizes from TGN/late endosomes to LDs. **A.** Oleate-loaded HeLa WT and SKO cells overexpressing CAV1-mCherry and GFP-RAB7A. **B.** Quantification of GFP-RAB7A localization to LDs. P = 0.0021, n = 15 cells for each condition from 3 replicates. **C.** Quantification of ARL8B Q75L recruitment to LDs in HeLa WT and SKO cells. P = 0.0048, n = 10 cells for each condition from 2 experiments. **D.** HeLa SKO cells overexpressing CAV1-mCherry and CFP-cdM6PR and loaded with oleic acid. **E.** Oleate-loaded HeLa WT and SKO cells overexpressing CAV1-GFP and RFP-VAMP7. **F.** Oleate-loaded HeLa SKO cells overexpressing CAV1-GFP. Chloroquine (Chlq.) is added to the medium during the oleic acid loading. **G.** Left: mean normalized CAV1-GFP signal around LDs per cell. Two-tailed Mann-Whitney analysis, P = 0.0014, n = 30 cells for KO and 15 for KO + Chlq. from 3 replicates. Right: mean CAV1-GFP PM signal (Fig. S5D). Two-tailed Mann-Whitney analysis, P = 0.8918, n = 15 for KO and 10 for KO + Chlq. cells from 3 experiments. **H.** Transmission electron microscopy of oleate-loaded HeLa WT and SKO cells overexpressing TGN38-HRP. Scale bar: 200 nm. **I.** Additional examples of LDs with TGN38-HRP-containing large structures near them. Scale bar: 500 nm. **J.** Top: fraction of LDs associated with TGN38-HRP vesicles per cell. P = 0.0171, n = 9 for WT and 7 for KO cells. Bottom: number of TGN-HRP-containing late endosomes. P = 0.0009, n = 9 for WT and 8 for KO cells. Unless indicated, scale bars are 5 µm.

To further test our hypothesis, we transfected cells with a CFP construct of the cation-dependent mannose-6-phosphate receptor (M6PR), which carries hydrolases from the Golgi to lysosomes. No M6PR accumulation was detected near LDs or colocalizing with CAV1 in WT cells. In contrast, M6PR vesicles were docked to LDs positive for CAV1 in SKO cells (Fig. 3D). These observations suggest that in SKO cells, the mislocalized CAV1 to LDs resulted from trafficking defects between the TGN and the PM, causing the rerouting of CAV1 through late endosomes and lysosomal pathways.

If CAV1 were delivered to LDs via TGN/late endosomal compartments, this would occur through hemifusion of the compartment outer monolayer with the LD monolayer^38,39^. This topology allows monotopic proteins to relocate from a bilayer to a monolayer, similar to the ER to LD protein trafficking pathway^40^. Therefore, we examined various SNARE proteins that could mediate such fusion (Fig. S5A-C). Out of many tested SNAREs, we found a clear colocalization of VAMP7 with CAV1 near LDs in SKO cells (Fig. 3E). We also sporadically detected Ykt6 dots near LDs (Fig. S5C), suggesting these SNARE components contribute to TGN/late endosome-LD fusion. Interestingly, these fusion machinery components are also involved in lysosome and autophagosome fusion^41^, which can be blocked by chloroquine^42^. Treating SKO cells with chloroquine showed a dramatic reduction in CAV1 relocation to LDs and the accumulation of vesicles near the LDs (Fig. 3F-G). These data strongly suggest that a controlled and direct physical contiguity between TGN/late endosome compartments and LDs is mediated by a fusion involving SNAREs.

Lastly, we used EM to provide evidence of such interactions. We transfected the cells with a horseradish peroxidase-TGN38 construct (TGN38-HRP), suitable for detecting TGN-derived compartments. We observed TGN38-containing vesicles and TGN38-positive late endosomes much more frequently in SKO than WT cells, and these structures were often close to LDs in SKO cells (Fig. 3H-J, S5E-G), consistent with our fluorescent imaging data. In a few instances, we captured interfacial contiguity between the HRP-TGN38-containing compartments and LDs (Fig. 3H, SKO), confirming the possibility of direct material transfer from the TGN/late endosomes to LDs. Based solely on this data, one might speculate direct physical contact between the TGN and LD delivers CAV1 to LDs. However, combining it with the localization of M6PR, RAB7, and lysosomal proteins to LDs supports the controlled and physical contiguity model between TGN/endolysosome compartments and LD.

### Excess ceramides shift CAV1 localization from PM to LDs

Given the importance of cholesterol and ceramide/SM in the sorting of TGN cargo and the impairment of CAV1 TGN-to-PM trafficking caused by the blockade of CERT or OSBP^43^, particularly CERT, we tested the impact of excess cholesterol or ceramide on CAV1 trafficking.

We exposed WT cells to excess cholesterol and induced a liquid crystalline phase in LDs^44^, indicating its storage as a cholesterol ester (Fig. S6A). In this condition, CAV1 was still present at the PM in WT cells and absent from LDs. Excess cholesterol did not phenocopy the impact of seipin deletion. In striking contrast, in WT cells supplemented with ceramides, overexpressed and endogenous CAV1 were redirected to LDs at the expense of PM localization (Fig. 4A-C, S6B), mimicking the SKO cells. This suggests that ceramide accumulation is related to the phenotype in the SKO cells. To track the trajectory of ceramides, we supplemented cells with ceramides and NBD-ceramides. In WT cells, the NBD signal was mainly localized to the LD interior, indicating its storage (Fig. 4D, S6C). This was concurrent with CAV1 relocation to LDs. In SKO cells, the NBD signal was also in the LD interior (Fig. 4D, S6C), but we observed events where it was recruited to the LD surface, colocalizing with CAV1 (Fig. S6D). These data suggest that cells store excess ceramides in LDs, likely as acylceramides^45^. However, part of the ceramides leaked to the Golgi (Fig. 4D) and ended up in late endosomes, probably in the form of SM or glycosylated ceramides, promoting CAV1 delivery to LDs.

**Figure 4:**
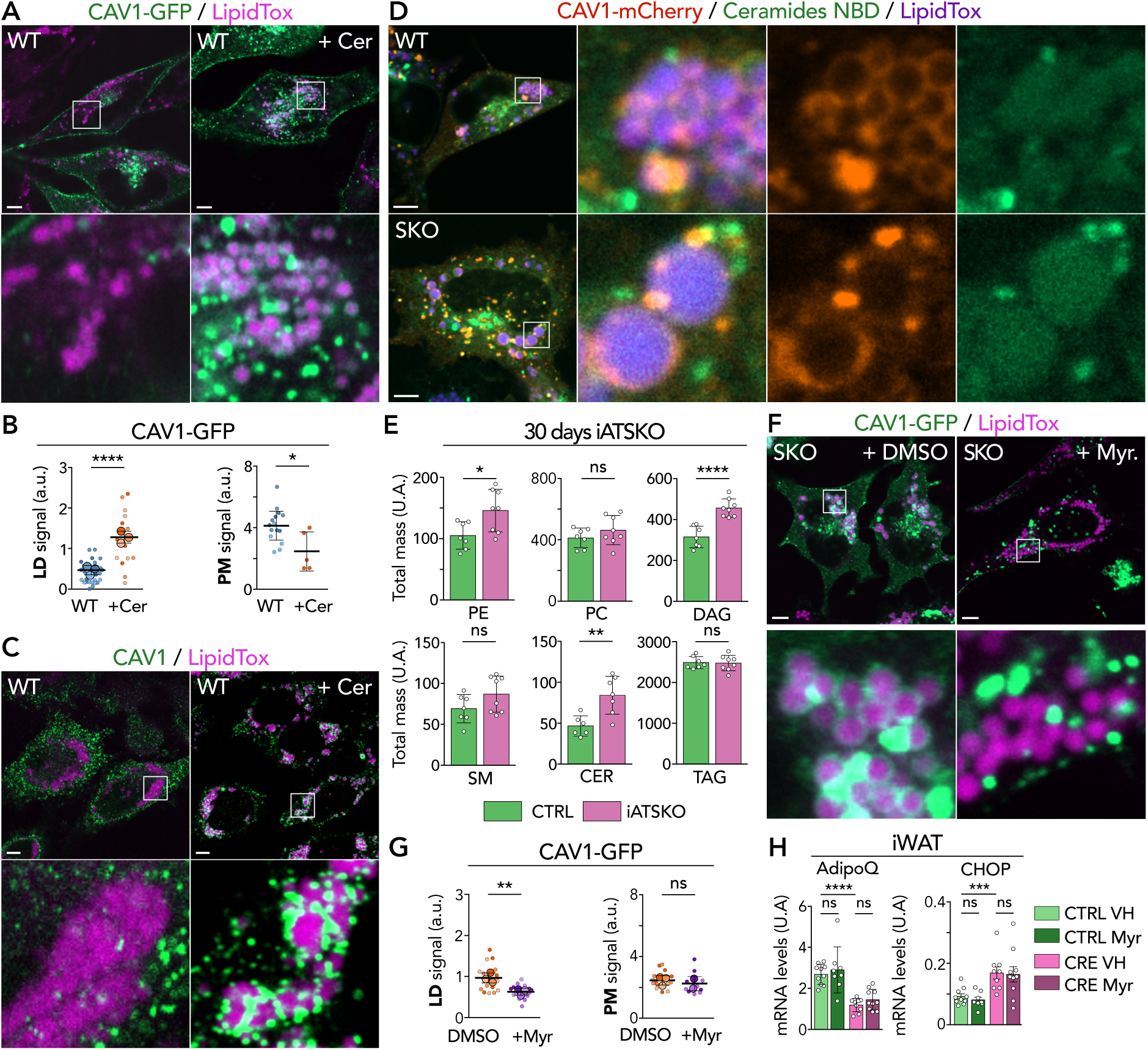
Excess ceramides shift CAV1 localization from PM to LDs. **A.** Oleate-loaded HeLa WT cells overexpressing CAV1-GFP and supplemented with 50 µM of ceramides (Cer.). **B.** Left: mean normalized CAV1-GFP signal around LDs per cell. P < 0.0001, n = 30 for WT and 15 for WT + Cer. cells from 3 experiments. Right: mean CAV1-mCherry PM signal. P = 0.0103, n = 15 for WT cells from 3 replicates and 5 for WT + Cer. **C.** Immunofluorescence of HeLa WT cells against the N-terminal part of CAV1 after incubation with oleic acid for 24 hours and 50 µM of ceramides. **D.** HeLa WT and SKO cells overexpressing CAV1-mCherry; 50 µM of ceramides containing 20% of ceramides-NBD is added to the medium during LDs formation by oleate addition. **E.** Whole tissue lipidomics of CTRL and iATSKO cells, 30 days after induction of the KO. PE, PC, DAG, SM, CER, and TAG have respective P-values of 0.0201, 0.2592, < 0.0001, 0.8836, 0.1088 and 0.0046; n = 7 for CTRL and 8 for iATSKO. **F.** Oleate-loaded HeLa SKO cells overexpressing CAV1-GFP are incubated with 2.5 µM of Myriocin (Myr) during the transfection and maintained afterward. A control is done with DMSO. **G.** Left: mean normalized CAV1-GFP signal around LDs per cell. P = 0.0019, n = 15 each from 3 replicates. Right: mean CAV1-GFP PM signal. P = 0.4216, n = 15 for KO + DMSO and 10 for KO + Myr. cells from 2 replicates. **H.** Transcriptomics analysis of AdipoQ and CHOP expression of inguinal iATSKO and control cells in the presence or not of myriocin. A two-tailed Mann-Whitney analysis is performed with respective P-values of 0.9654, 0.4967, < 0.0001, 0.4082, 0.7197, and 0.0006. n for AdipoQ experiments is 10 for CTRL VH, 9 for CTRL Myr, 9 for CRE VH, and 10 for CRE Myr cells; n for CHOP experiments is 10 for CTRL VH, 8 for CTRL Myr, 9 for CRE VH and 10 for CRE Myr. Scale bars: 5 µm.

We analyzed the lipid composition of the AT from CTRL and iATSKO mice, 14 and 30 days of knockout induction. The lipidome profile was altered in many species upon seipin depletion. Notably, phosphatidylethanolamine (PE), phosphatidylcholine (PC), SM, and ceramide levels increased. Triacylglycerol and diacylglycerol levels were similar despite a tendency for diacylglycerol to increase (Fig. 4E; S6E). Interestingly, weight correlation network analysis of lipidomic and transcriptomic datasets revealed that in iATSKO AT, the ceramides levels were positively correlated with ER stress markers and negatively associated with the mRNA levels of the desaturase SCD1 (Fig. S7A-B). Ceramides are made in the ER, so we purified the ER from these cells. We found that the iATSKO condition significantly increased diacylglycerol and ceramide levels, while other significant lipids did not change substantially (Fig. S6F). These results indicate the formation of excess ceramides when seipin is removed.

To reduce ceramides, we treated SKO cells with myriocin to block serine palmitoyltransferase (SPT), the rate-limiting enzyme of ceramide synthesis. This abolished the localization of CAV1 to LDs, and CAV1-containing vesicles remained near the LDs (Fig.4F-G). However, PM localization was not restored as in WT (Fig. 4G, S6G), likely because the inactivation of SPT significantly reduced ceramides and complex sphingolipids, which are essential for the TGN and many other cellular functions.

We repeated the experiment by treating iATSKO mice with myriocin. The adipose tissue of iATSKO displays a marked dysfunction characterized by a decreased expression of mature adipocyte markers such as AdipoQ and the induction of ER stress markers such as the pro-apoptotic transcription factor CHOP. Whereas myriocin successfully lowered the ceramide levels in the adipose tissue of iATSKO mice, it did not increase the mRNA level of AdipoQ nor reduce the expression of CHOP (Fig. 4H). Therefore, in this context, solely decreasing ceramides was insufficient to tackle the adipose tissue failure induced by seipin deficiency, consistent with the lack of rescue of CAV1 PM localization by SPT inhibition (Fig. 4F-G).

### Regulation of the secretory pathway by the glycerolipid-ceramide branch point

Ceramides originate from saturated fatty acids such as palmitate or stearate^46^. We did a lipidomics analysis of the Golgi-ER fraction and found increased lipid saturation in ER fractions of SKO cells relative to WT cells (Fig. 5A). Palmitate is a substrate for Acyl-CoA Synthetase, which activates it for ceramide synthesis. This pathway competes with palmitate elongation by Elovl6 and desaturation by SCD1 for incorporation into glycerolipids^47^ (Fig. 5B). Interestingly, in iATSKO AT, the mRNA levels of Elovl6 and SCD1 were over time inversely correlated with ceramides levels (Fig. S7A-B). Proteins involved in palmitic acid consumption were increased, while those involved in synthesis or import were decreased (Fig. S7C), suggesting a strategy of the cells to reduce saturated lipids’ accumulation.

**Figure 5:**
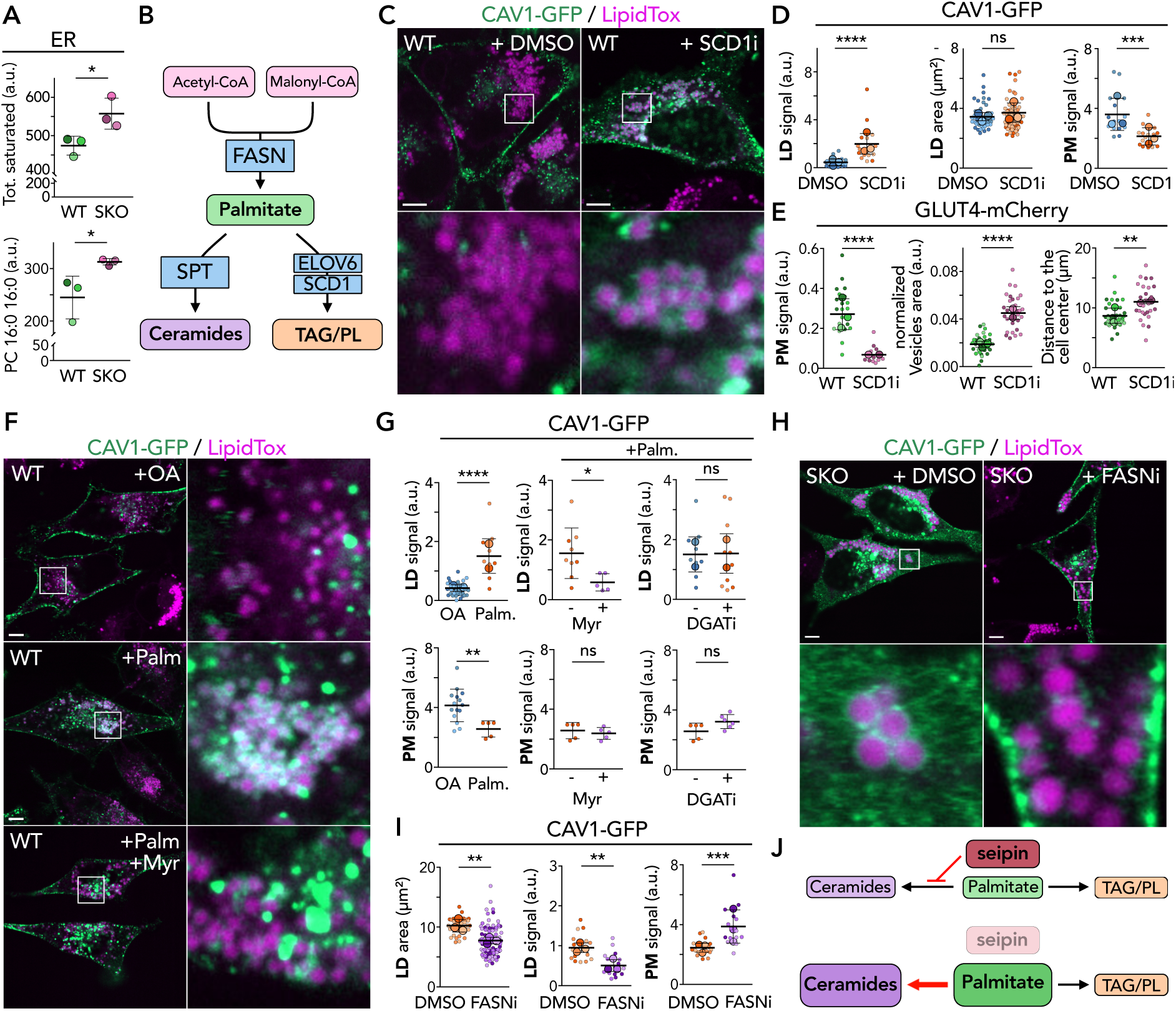
Regulation of the secretory pathway by the glycerolipid-ceramide branch point. **A.** Top: comparison of total saturated lipid content from ER fraction lipidomics analysis between WT and SKO. P = 0.0382, n = 3 experiments. Bottom: normalized PC 16:0 16:0 count from ER fraction. P = 0.0462; 3 experiments. **B.** Schematic representation of palmitic acid metabolic pathways and the associated enzymes and metabolites. **C.** Oleate-loaded HeLa WT cells overexpressing CAV1-GFP. 1 µM of an SCD1 inhibitor (SCD1i) is added 5 hours before transfection and maintained afterward. A control is done with DMSO. **D.** Left and Middle: mean LD area and normalized CAV1-GFP signal around LDs per cell. A two-tailed Mann-Whitney analysis is performed. LDs’ area: P = 0.1552, n = 54 for WT cells and 89 for KO cells from 3 experiments; CAV1-GFP signal around LDs: P< 0.0001, n = 21 WT + DMSO and 15 WT + SCD1i cells from 3 replicates. Right: mean CAV1-GFP PM signal normalized by basal cell signal. Two-tailed Mann-Whitney analysis, P = 0.0003, n = 15 cells each from 3 replicates. **E.** Quantification of GLUT4-mCherry overexpressing cells after SCD1 inhibition. From left to right: GLUT4-mCherry PM mean signal normalized by mean vesicle signal (P < 0.0001, n = 21 for WT and 13 for WT + SCD1i cells from 3 and 2 replicates); the sum of vesicles’ area divided by cell area (P < 0.0001, n = 35 for WT and 28 for WT + SCD1i cells from 3 and 2 replicates); GLUT4-mCherry vesicles’ dispersion in the cell (two-tailed Mann-Whitney analysis: P = 0.0011, n = 34 for WT and 26 for WT + SCD1i cells from 3 and 2 experiments, respectively). **F.** HeLa WT cells overexpressing CAV1-GFP, incubated with 2.5 µM of Myriocin (Myr) or 2.5 µM DGAT1/2 inhibitors (DGATi). After transfection, a palmitate-oleate mixture is added for 24 hours. In DGATi, Oleic acid induced LDs first; the inhibitors were added 5 hours before palmitate-oleate addition. **G.** Top: mean normalized CAV1-GFP signal around LDs per cell. Top left: quantification when oleic acid was alone or mixed with palmitate. P < 0.0001, n = 29 for WT and 9 WT + palmitate-oleate cells from 3 and 2 replicates. Top middle: quantification of WT cells loaded with a palmitate-oleate mix with or without Myriocin (Myr). P = 0.0303, n = 9 WT – Myr and 5 WT + Myr cells. Top right: quantification of WT cells loaded with oleic acid to induce LDs and then incubated with a mix of palmitate-oleate with or without 2.5 µM of DGAT1/2 inhibitors. P = 0.9701, n = 9 for WT and 10 for WT + DGATi cells from 2 replicates. Bottom: mean CAV1-GFP PM signal. Bottom left: quantification when oleate is alone or mixed with palmitate. Two-tailed Mann-Whitney analysis: P = 0.0037, n = 15 for WT and 5 for WT + Palm cells. Bottom middle: quantification of WT cells loaded with a palmitate-oleate mix in the presence or absence of Myriocin (Myr). Two-tailed Mann-Whitney analysis, P = 0.4206, n = 5 cells for each condition. Bottom right: quantification of WT cells incubated with a mix of palmitate-oleate in the presence or not of 2.5 µM of DGAT1/2 inhibitors. Two-tailed Mann-Whitney analysis: P = 0.1255, n = 5 for WT and 6 for WT + DGATi cells. **H.** Oleate-loaded HeLa WT cells overexpressing CAV1-GFP. 1µM of a FASN inhibitor (FASNi) is added 5 hours before transfection and maintained afterward. Loading control is with DMSO. **I.** Left: mean LD area and mean normalized CAV1-GFP signal around LDs per cell. A two-tailed Mann-Whitney analysis is done for the LD area’s quantification, P < 0.0001, n = 34 for KO + DMSO cells and 79 for KO + FASNi cells from 3 replicates. Middle: for CAV1-GFP recruitment to LDs, P = 0.0010, n = 15 cells each from 3 experiments. Right: mean CAV1-GFP PM signal. P = 0.0010, n = 15 cells each from 3 experiments. **J.** Model for seipin impact on ceramides and glycerolipids. Scale bars: 5 µm.

We tested whether directing palmitate toward ceramides or glycerolipids could regulate CAV1 localization to the PM and LDs. We inhibited SCD1 (SCD1i) in WT cells to promote palmitate/stearate accumulation and loaded them with oleate to induce LD formation. Despite the presence of seipin, this maneuver significantly reduced the localization of both overexpressed and endogenous CAV1 to the PM and strongly promoted its relocation to LDs (Fig. 5C-D, S7F-G). The size of the LDs was not significantly impacted. Concurrently, SCD1 inhibition strongly impaired the translocation of GLUT4 vesicles to the PM (Fig. 5E, S8A), caused their dispersion, and affected the localization of SNAP23, CD36, OSBP and P4M (Fig. S8B-I). SCD1i mimicked all the identified phenotypes of seipin KO in WT cells, suggesting that the two proteins work in parallel^48,49^. Accordingly, the overexpression of SCD1 in SKO cells was strikingly efficient and sufficient to rescue CAV1 PM localization and prevent its binding to LDs in SKO cells (Fig. S7H-I).

The above data suggested that the overaccumulation of palmitate led to its channeling into ceramides, which caused the observed CAV1 phenotypes. This implies that fluctuations in palmitate levels and, therefore, in ceramide levels modulate CAV1 trafficking, organelle homeostasis, and the secretory pathway. To test this, we supplemented WT cells with palmitic acid and found that it was sufficient to relocate CAV1 from the PM to LDs (Fig. 5F). Providing myriocin in this condition to block ceramide formation from the delivered palmitate prevented CAV1 LD localization. As previously, the protein accumulated in vesicles close to LDs, similar to myriocin-treated SKO cells. Blocking diacylglycerol O-acyltransferase (DGAT) enzymes to prevent the incorporation of the palmitate or ceramide into LDs did not enhance the phenotype, indicating that the accumulating ceramides in membranes, not acylceramides fabricated by DGAT2^45^ and stored in LDs, was responsible for CAV1 localization to LDs (Fig. 5F-G, S8J). Lastly, since palmitate levels were elevated in SKO cells, we pharmacologically blocked the activity of FASN, the critical enzyme mediating fatty acid synthesis (Fig. 5B). This nicely restored CAV1 PM localization and abolished its LD signal (Fig. 5H, S8K). These data indicated that the funneling of palmitate toward ceramide versus glycerolipid synthesis controls organelle dynamics and protein trafficking.

## Discussion

Seipin regulates LD nucleation and growth, but indirect observations suggest it has a broader role in lipid metabolism^50–52^. While LDs can form without the involvement of proteins^53^, seipin coordinates this process to ensure they form and function correctly. When seipin is deficient, adipogenesis is impaired, and when it is removed, adipose tissues are lost even though LDs may still be present. In contrast, mice lacking DGAT enzymes develop adipose tissues without LDs^54^. These findings decorrelate the involvement of seipin in the physical control of LD formation, lipid metabolism, and adipogenesis, highlighting its complex and multifaceted functions.

Here, we discovered that seipin deficiency causes the accumulation of palmitate, which leads to an increase in ceramide levels. This accumulation disrupts the function of TGN, impairing its ability to sort and transport cargo properly. As a result, CAV1 cannot be adequately trafficked to the PM, thereby reducing caveolae formation. Instead of targeting the PM, CAV1 is rerouted through the endosomal maturation system. It relocates to LDs by the fusion of TGN/late endosome-LD with LDs in a ceramide-dependent manner. When we blocked ceramide synthesis, the endosome-LD fusion was prevented, but this did not restore the normal relocalization of CAV1 to the PM. However, when we overexpressed SCD1 or inhibited FASN, we observed a rescue of both the LD and PM phenotypes. This indicates that the abnormal accumulation of palmitate is the primary cause of the defects observed in CAV1 localizaiton. Supplementation with palmitate or ceramides to WT cells reproduced the identical seipin deficiency phenotypes, confirming their role in this process. Overall, our findings underscore the pivotal role of seipin in regulating protein trafficking and organelle dynamics. Seipin functions at the crossroads of ceramide and glycerolipid biosynthesis, modulating the allocation of synthesized palmitate to these lipid species. (Fig. 5J).

Seipin assembles into an oligomeric, donut-shaped structure consisting of 11-12 monomers^55–57^, providing multiple potential interaction sites that could help coordinate local metabolic reactions. Indeed, interestingly, seipin interacts with several enzymes involved in the synthesis of ceramides and glycerolipids, such as SPT, SCD1, glycerol-3-phosphate acyltransferases, DGAT2, 1-acylglycerol-3-phosphate O-acyltransferase 2 (AGPAT2), and Lipin1^49–51,58,59^. These interactions suggest that seipin may regulate these enzymes based on metabolic signals and coordinate lipid-related biochemical reactions in specific regions^60^. For example, seipin is recruited to the mitochondria-associated membrane (MAM) during lipogenesis, where some LDs form and others are supplied with lipids^61^. This recruitment aligns with observations of SCD1 and DGAT2 accumulation at ER-mitochondria contact sites^58^, indicating that these enzymes may need to be concentrated at such locations in a seipin-dependent manner^49^. In the absence of seipin, overproducing enzymes like SCD1 may achieve the necessary local concentration, which likely explains why SCD1 overexpression rescued CAV1 in the SKO cells. Based on these findings, we propose that specific seipin complexes, potentially those at MAMs, create an environment balancing palmitate incorporation into glycerolipids and LDs versus ceramides. This is achieved by efficiently routing lipid substrates and products to enzymes near the seipin complex, ensuring controlled and efficient lipid metabolic processes.

The influence of seipin on the ceramide/glycerolipid balance could be determined by the rate of palmitate synthesis or uptake, which is governed by the cell’s metabolic demands and seipin levels. When ceramide levels reach a critical threshold, cargo transport from the TGN to the PM is impaired, particularly for those that rely on membrane order. This disruption can cause the mislocalization of cargos like CAV1 to LDs via the endolysosomal system, potentially serving as a cellular strategy to alleviate stress caused by lipid homeostasis disturbances. This pathway might be activated in specific cell types under particular physiological or pathophysiological conditions and could be constitutively active in the absence of seipin. In cells with low seipin levels, such as hepatocytes, this pathway may be more frequently triggered to deliver endocytosed lipids to LDs^31^. This represents a novel mechanism by which physical contiguity between endosomes and LDs regulates lipid metabolism^39^.

Our findings show that seipin deficiency leads to the mislocalization of CAV1 to the PM and the decrease in caveolae. Similarly, CAV1 and Cavin1 deficiency are responsible for CGL3 and CGL4, respectively, decreasing caveolae. Lastly, recently, in newborn mice deficient for AGPAT2, the gene whose mutation causes CGL1, the adipocyte displays an early caveolae loss that precedes lipodystrophy^62^. Therefore, all CGLs have the loss of caveolae in common, which could be a key element in the primary adipocyte dysfunction that characterizes CGLs. Notably, several pieces of evidence produced in CAV1-deficient adipocytes support the crucial role of caveolae in lipid handling and insulin signaling in the adipocytes^63^.

CAV1 plays a crucial role in maintaining lipid raft domains, which are essential for the proper localization and stabilization of the insulin receptor (IR) at the PM in adipocytes. In addition to supporting IR function, CAV1 regulates downstream signaling pathways, including the secretion of adipokines and free fatty acids, vital for normal adipocyte function and lipid storage. When CAV1 is dysfunctional, it accelerates the degradation of both IR and GLUT4^14,24^, leading to impaired insulin signaling and systemic insulin resistance. This dysfunction is strongly linked to metabolic disorders, such as type 2 diabetes and obesity. Given seipin’s role in CAV1 mislocalization to the PM, our study uncovers a cell-autonomous mechanism where seipin deficiency promotes insulin resistance through the accumulation of ceramides.

Finally, our data suggest that the accumulation of ceramide, palmitate, and stearate, all known predictors of incident type 2 diabetes^64^, could also be associated with CAV1, mislocalization, and, thus, IR and GLUT4 cycling. This opens a new avenue for pathophysiological studies of human insulin resistance, possibly leading to innovative pharmacological interventions.

## Notes

### Competing Interest Statement

The authors have declared no competing interest.

