## Supplementary materials for "Seipin Regulates Caveolin-1 Trafficking and Organelle Crosstalk"

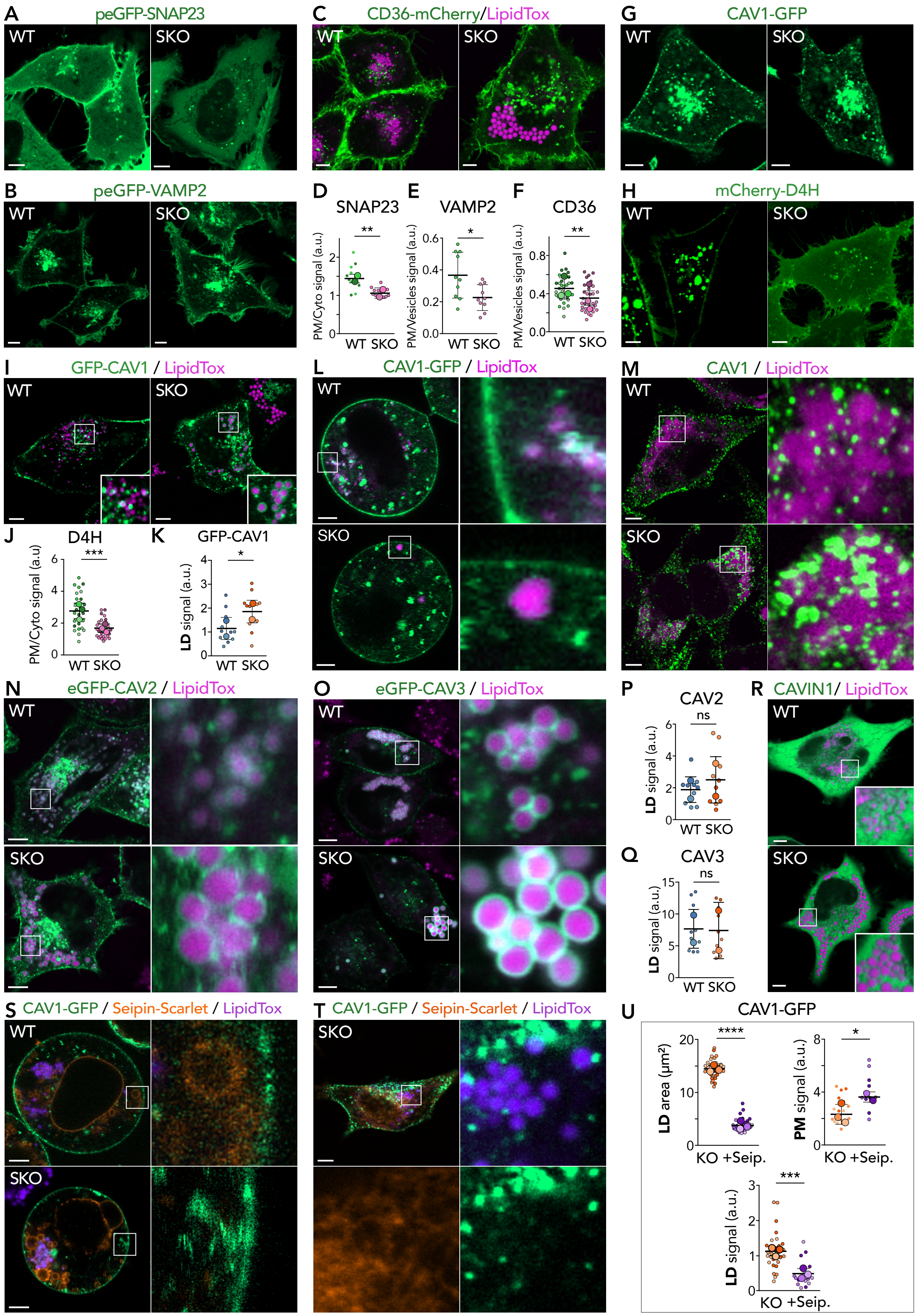


**Figure S1: A, B.** HeLa WT and SKO cells overexpressing p-eGFP-SNAP23 or peGFP-VAMP2. **C.** Oleate-loaded HeLa WT and SKO cells overexpressing CD36-mCherry. **D, E.** Normalized PM signal of SNAP23 and VAMP2. Panel D: P = 0.034, n = 10 for WT and KO cells from 2 replicates. Panel E: P = 0.0153, n = 10 for WT and KO cells. **F.** CD36-mCherry mean PM signal normalized by the mean signal of vesicles. P = 0.0079, n = 30 for WT and KO from 3 replicates. **G.** HeLa WT and SKO cells overexpressing CAV1-GFP. **H.** HeLa WT and SKO cells overexpressing mCherry-D4H. **I.** Oleate-loaded HeLa WT and SKO cells overexpressing an N-terminal GFP tagged CAV1 (GFP-CAV1). **J.** Normalized mCherry-D4H mean PM signal. P = 0.0002, n = 24 for WT and KO cells from 3 experiments. **K.** Normalized eGFP-CAV1 mean signal around LDs per cell. P = 0.0309, n = 11 for WT and KO cells from 2 replicates. **L.** Oleate-loaded and swollen HeLa WT and SKO cells overexpressing CAV1-GFP. **M.** Immunofluorescence against N-terminal part of CAV1 in oleate-loaded HeLa WT and SKO cells. **N, O.** Oleate-loaded HeLa WT and SKO cells overexpressing eGFP-CAV2 or eGFP-CAV3. **P, Q.** Normalized eGFP-CAV2 or eGFP-CAV3 mean signal around LDs. Panel P: P = 0.3256, n = 10 for WT and KO cells from 2 replicates. Panel Q: P = 0.8774, n = 10 for WT and KO cells from 2 replicates. **R.** HeLa WT and SKO cells overexpressing CAVIN1 and oleate-loaded. **S.** Oleate-loaded and swollen HeLa WT and SKO cells overexpressing CAV1-GFP and seipin-mScarlet. CAV1 PM localization is rescued. **T.** Oleate-loaded HeLa SKO cells overexpressing CAV1-GFP and seipin-mScarlet. CAV1 is excluded from LDs. **U.** Left and middle: mean LD area per cell and mean normalized CAV1-GFP signal around LDs per cell. A two-tailed Mann-Whitney analysis is done for LD's area (P < 0.0001 n =39 for KO and 35 for KO + Seipin cells from 3 replicates) and CAV1-GFP signal around LD (P=0.0001, n= 29 for KO and 15 for KO + Seipin cells from 3 experiments). Right: mean CAV1-GFP PM signal per cell normalized by a basal signal from experiments in C. P=0.0137, n= 15 for KO and 10 for KO + seipin cells, respectively 3 and 2 replicates. Scale bars: 5 µm.

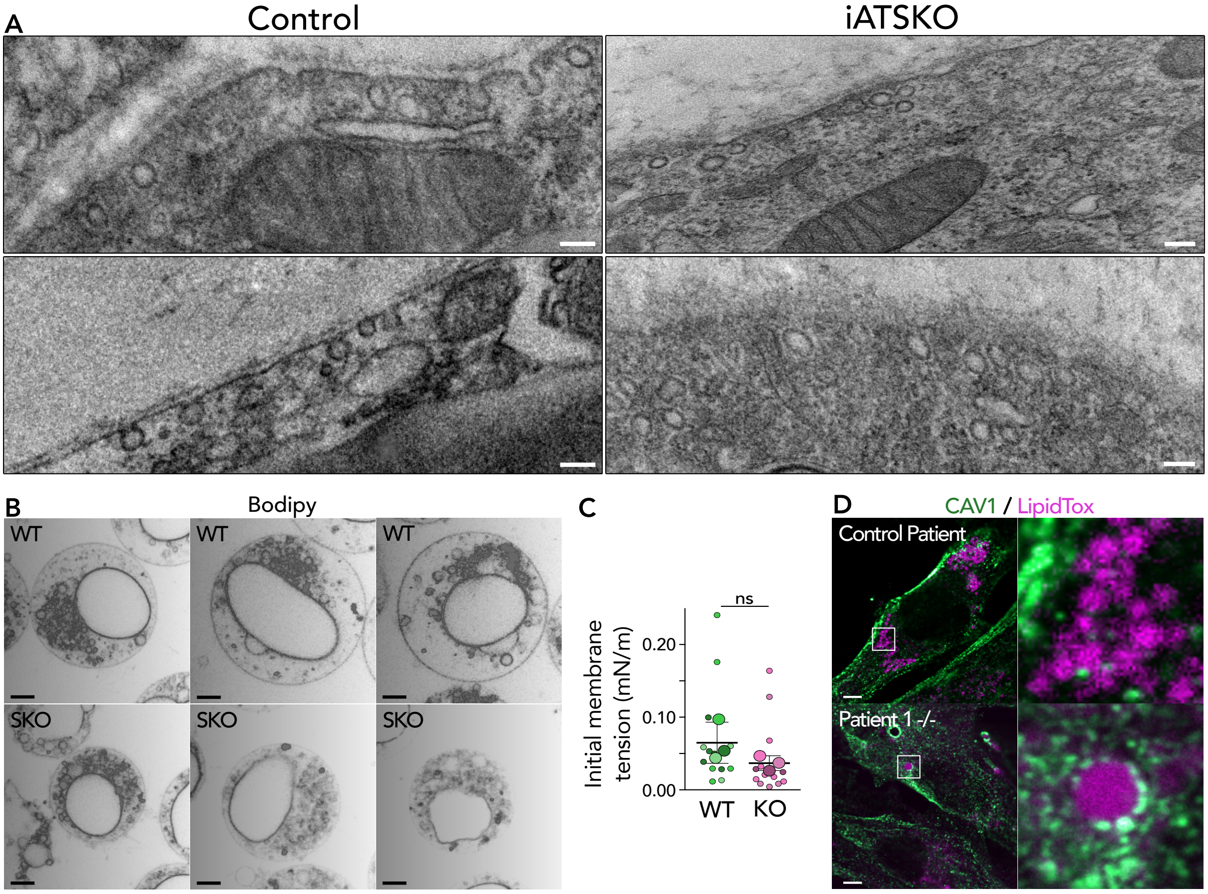


**Figure S2: A.** Additional transmission electron microscopy cell images from control and iATSKO mice. Scale bar: 100 nm**. B.** Additional cell images of swollen HeLa WT and SKO cells labeled with Bodipy. Scale bar: 5 µm. **C.** Quantification of the initial membrane tension from swollen HeLa WT and SKO cells labeled with Bodipy. Two-tailed Mann-Whitney analysis, P = 0.0723. n = 13 for WT and SKO cells from 3 replicates. **D.** Immunofluorescence against N-terminal part of CAV1 in fibroblasts from control and CGL2 patients. LDs are induced by oleate supply. Scale bar: 5 µm.

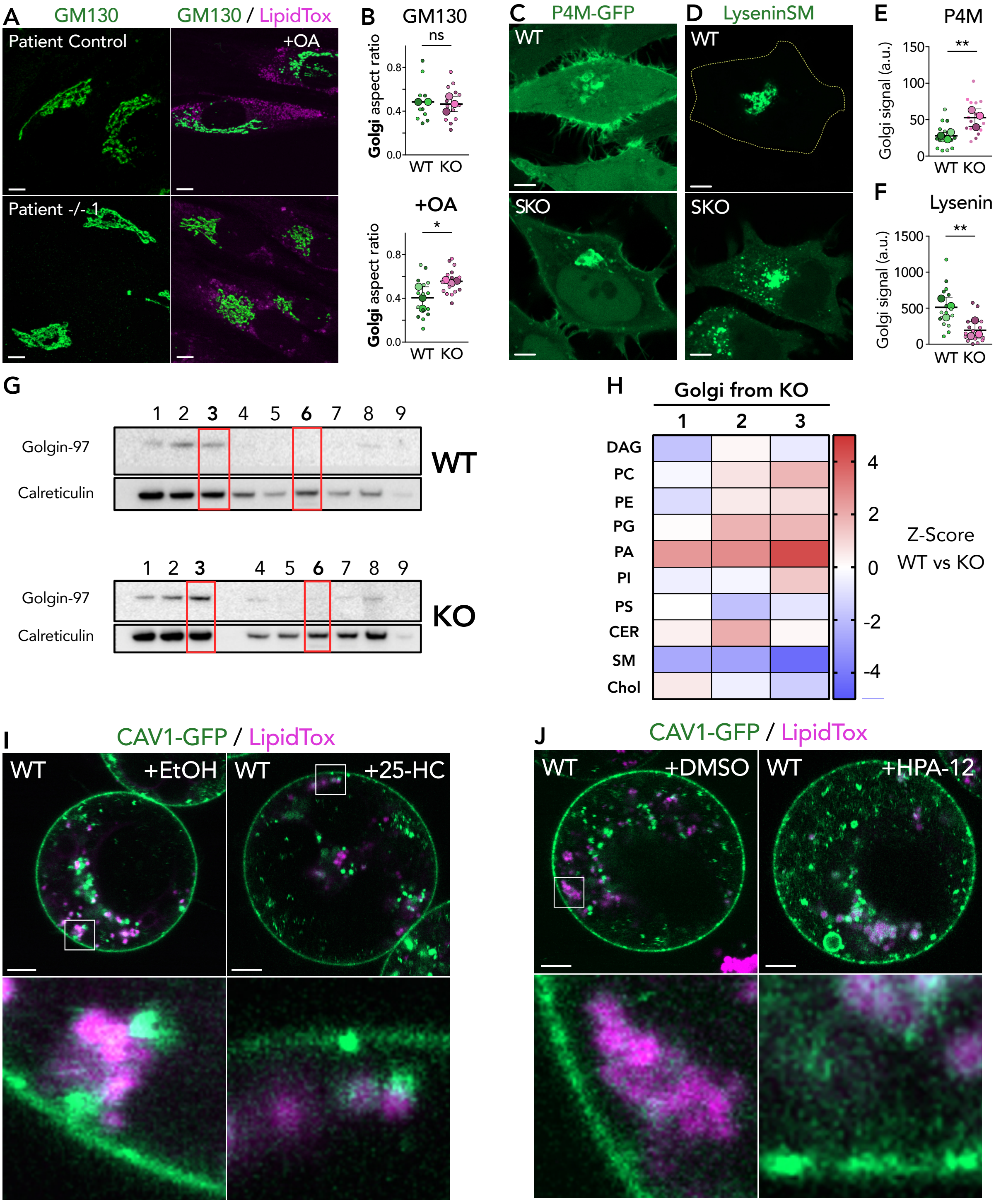


**Figure S3: A.** Immunofluorescence against Golgi resident protein GM130 in fibroblasts from control and CGL2 patients, after LD induction or not with oleic acid. **B.** Scatter plots of Golgi aspect ratio from the experiment in A. Top: without lipid loading, P = 0.7842, n = 10 control and 15 CGL2 patient cells, from respectively 2 and 3 replicates. Bottom: after LD induction, P = 0.0108, n = 15 for control and CGL2 patient cells, from 3 replicates. **C, D.** HeLa WT and SKO cells overexpressing the P4M-GFP PI4P or Lysenin sphingomyelin probes. **E, F.** Quantification of experiments in C and D. Panel E: P = 0.024, n = 15 for WT and KO cells, from 3 experiments. Panel F: P = 0.0011, n = 15 for WT and 15 for KO cells, from 3 replicates. **G.** Western blots of HeLa WT and SKO cell lysates separated by ultracentrifugation. The fractions enriched with the TGN are labeled using an antibody against Golgin-97, while the ER-corresponding fractions are labeled using an antibody targeting Calreticulin. Fractions 3 and 6 were chosen as Golgi-enriched fractions**. H.** Heatmap analysis of lipidomics data of

Golgi fractions. **I, J.** HeLa WT and SKO cells overexpressing CAV1-GFP, oleate-loaded and swollen. OSBP and CERT are inhibited with 2.5 μM of 25-HC or HPA-12. Scale bar: 5 μm.

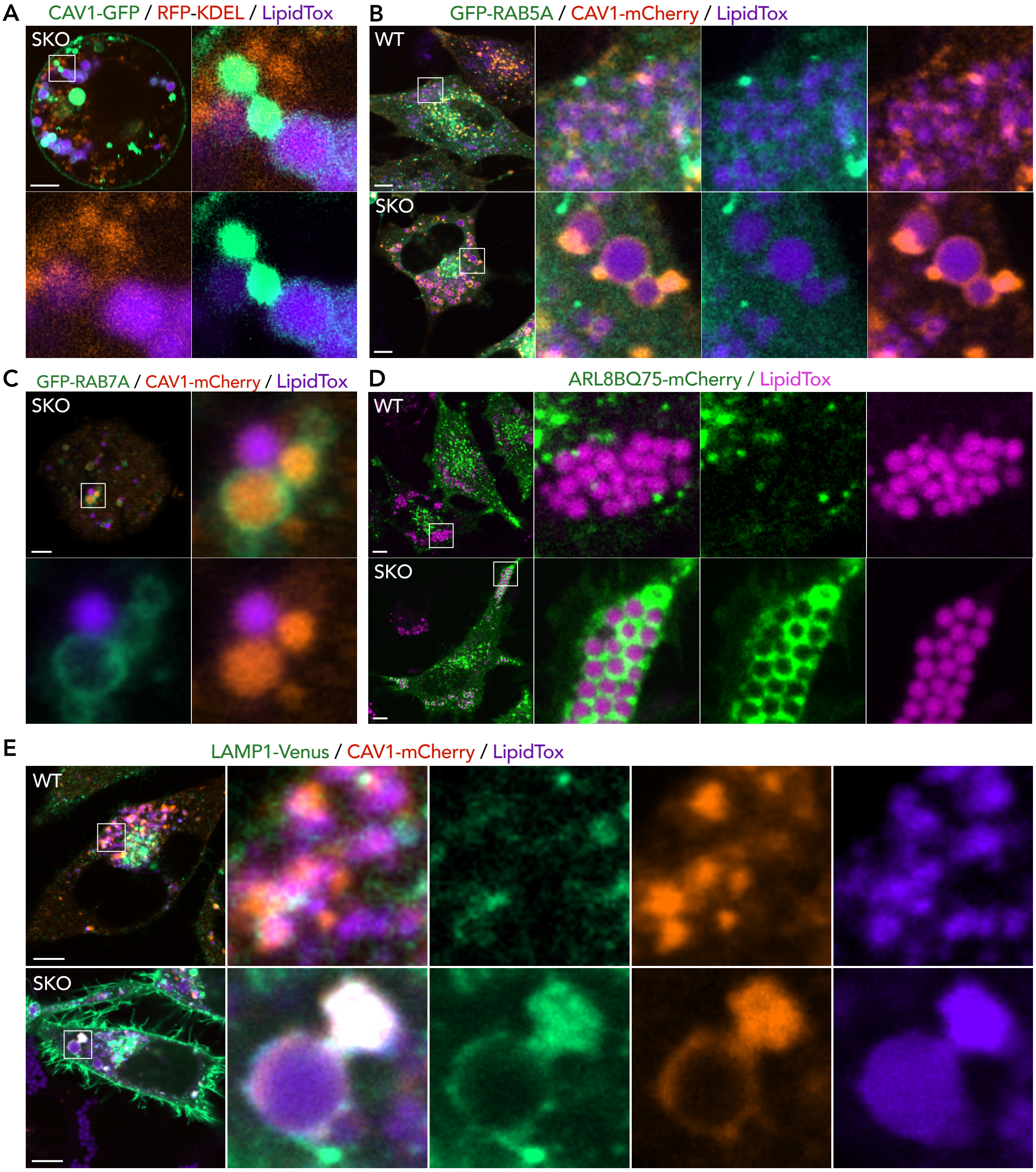


**Figure S4: A**. Oleate-loaded and swollen HeLa SKO cells overexpressing CAV1-mCherry and ER marker RFP-KDEL. **B.** Oleate-loaded HeLa WT and SKO cells overexpressing CAV1-mCherry and GFP-RAB5A. **C.** Oleate-loaded and swollen HeLa SKO cells overexpressing CAV1-mCherry and GFP-RAB7A. **D.** Oleate-loaded HeLa WT and SKO overexpressing ARL8B-Q75L-mCherry. **E.** Oleate-loaded HeLa WT and SKO overexpressing CAV1-mCherry and LAMP1-Venus. Scale bars: 5 μm.

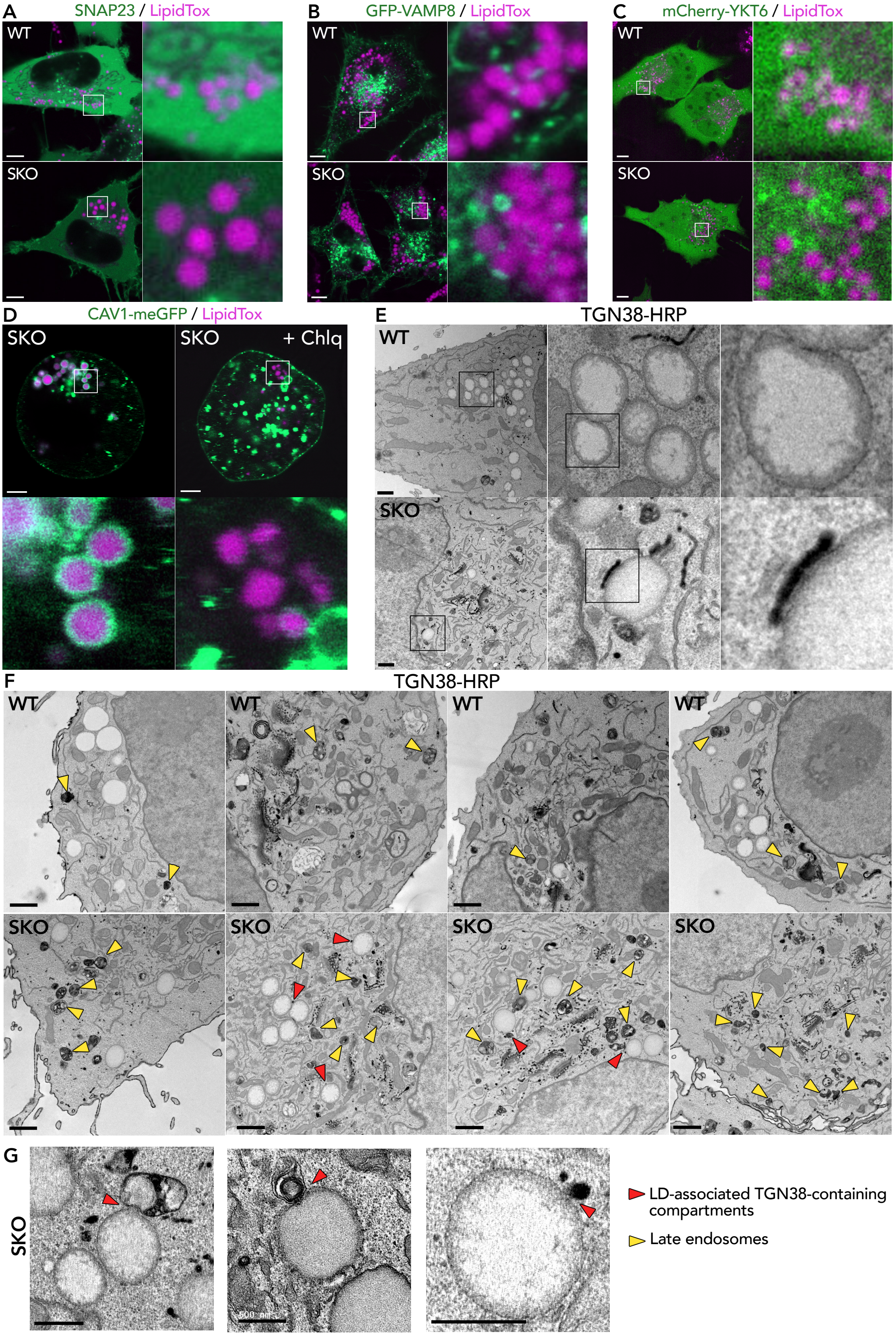


**Figure S5: A, B, C.** Oleate-loaded HeLa WT and SKO cells overexpressing pEGFP-SNAP23, GFP-VAMP8, or mCherry-YKT6. Scale bars 5 µm. **D**. Swollen HeLa SKO overexpressing CAV1-GFP, in the presence or not of chloroquine (Chlq.). Scale bar 5 µm. **E, F, G.** Transmission electronic microscopy of HeLa WT and SKO cells overexpressing TGN38-HRP and loaded with oleate (selected zoom-ins are shown in G). Yellow arrows correspond to late endosomes, and red ones to TGN38-containing compartments associated with LDs. Scale bars 100 nm in F and 500 nm in G.

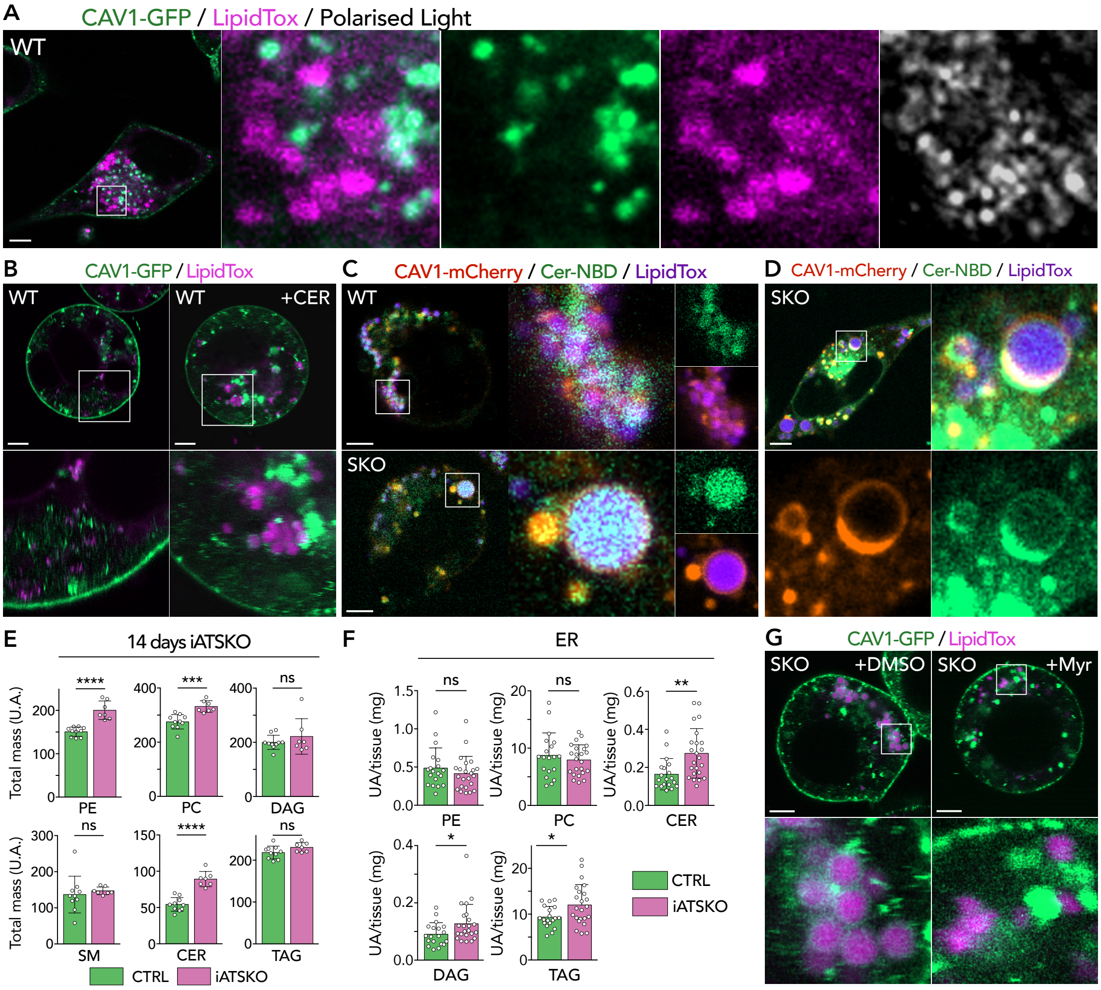


**Figure S6. A.** Cholesterol-fed WT HeLa cells transfected with CAV1-GFP. The polarized light signal indicates the accumulation of cholesterol ester in LDs. CAV1 signal is in vesicles close to LDs but not around LDs. **B, C.** Oleate-loaded and swollen HeLa WT and SKO cells overexpressing CAV1-GFP or CAV1-mCherry. 50 µM of ceramides is added to the media during LD formation. In panel C, 20% ceramide-NBD is added to the ceramide solution. **D**. Oleate-loaded HeLa WT and SKO cells overexpressing CAV1-mCherry. 50 µM of ceramides containing 20% ceramides-NBD are added to the medium. **E**. Whole tissue lipidomics of CTRL and iATSKO cells, 14 days after induction of the KO. PE, PC, CER, and TAG have P-values of < 0.0001, 0.0003, < 0.0001, and 0.0922. Two-tailed Mann-Whitney analysis for DAG and SM with respective P-values of 0.6691 and 0.1932. n = 10 for CTRL and 8 for iATSKO mice. **F**. ER extracted lipidomics analysis of CTRL and iATSKO cells. Two-tailed Mann-Whitney analysis for PE, PC, CER, DAG, and TAG with P-values of 0.3406, 0.7451, 0.0015, 0.0287, 0.0451. n = 19 for CTRL and 24 for iATSKO mice, from 2 replicates **G.** Oleate-loaded and swollen HeLa SKO cells overexpressing CAV1-GFP. Cells are incubated with 2.5 µM of Myriocin (Myr) during the transfection and maintained afterward. Control is with DMSO. Scale bar: 5 µm.

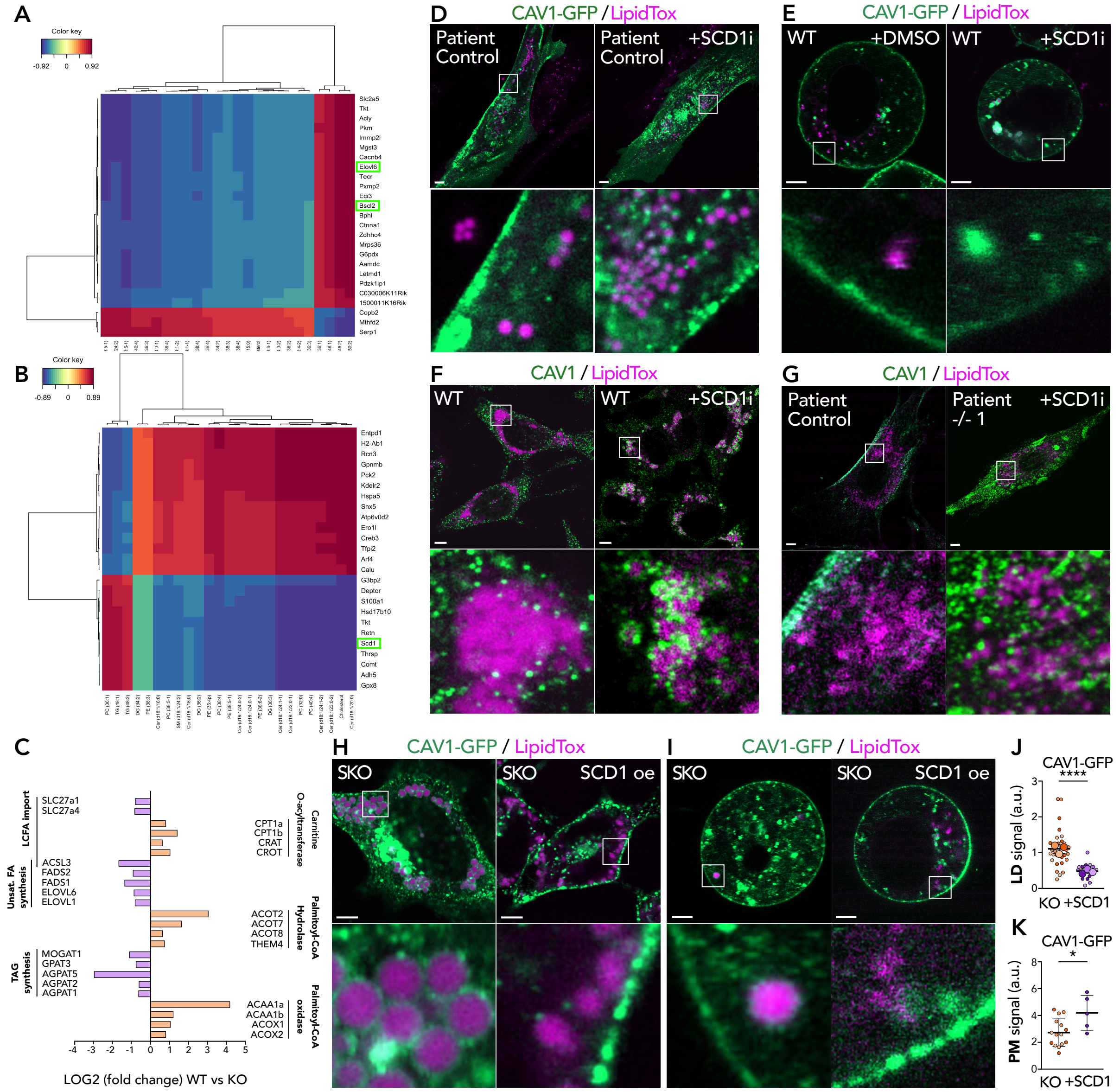


**Figure S7: A, B.** Heatmaps of correlation between mRNA levels and lipids species in iATSKO vs CTLR mice, 14 days and 30 days after KO. A green box highlights seipin, Evolv6, and SCD1. **C.** Proteomic analysis of ER extracts from CTRL and iATSO mice. **D.** Fibroblast from control patient overexpressing CAV1-GFP. 1 µM of SCD1 inhibitor (SCD1i) is added 5 hours before transfection and maintained afterward. LDs are induced by oleic acid supply. **E.** Oleate-loaded and swollen HeLa WT L cells overexpressing CAV1-GFP. 1 µM of SCD1 inhibitor (SCD1i) is added 5 hours before transfection and maintained afterward. Control is with DMSO. **F, G**. Immunofluorescence against N-terminal part of CAV1 in HeLa WT cells and fibroblast from control patient. 1 µM of SCD1i is added 5 hours before inducing LDs. **H**. Oleate-loaded HeLa SKO cells overexpressing CAV1-GFP and SCD1. **I**. Oleate-loaded and swollen HeLa SKO cells overexpressing CAV1-GFP and SCD1. **J**. Mean normalized CAV1-GFPsignal around LDs per cell, from experiments in H. Two-tailed Mann-Whitney analysis, $P < 0.0001$. $n = 29$ for KO and 15 for KO + SCD1 cells, from 3 replicates. **K**. Mean CAV1-GFP PM signal normalized by the basal cell signal from experiments shown in I. $P = 0.0176$. $n = 15$ for KO and 5 for KO + SCD1 from 1 experiment. Scale bars: 5 µm.

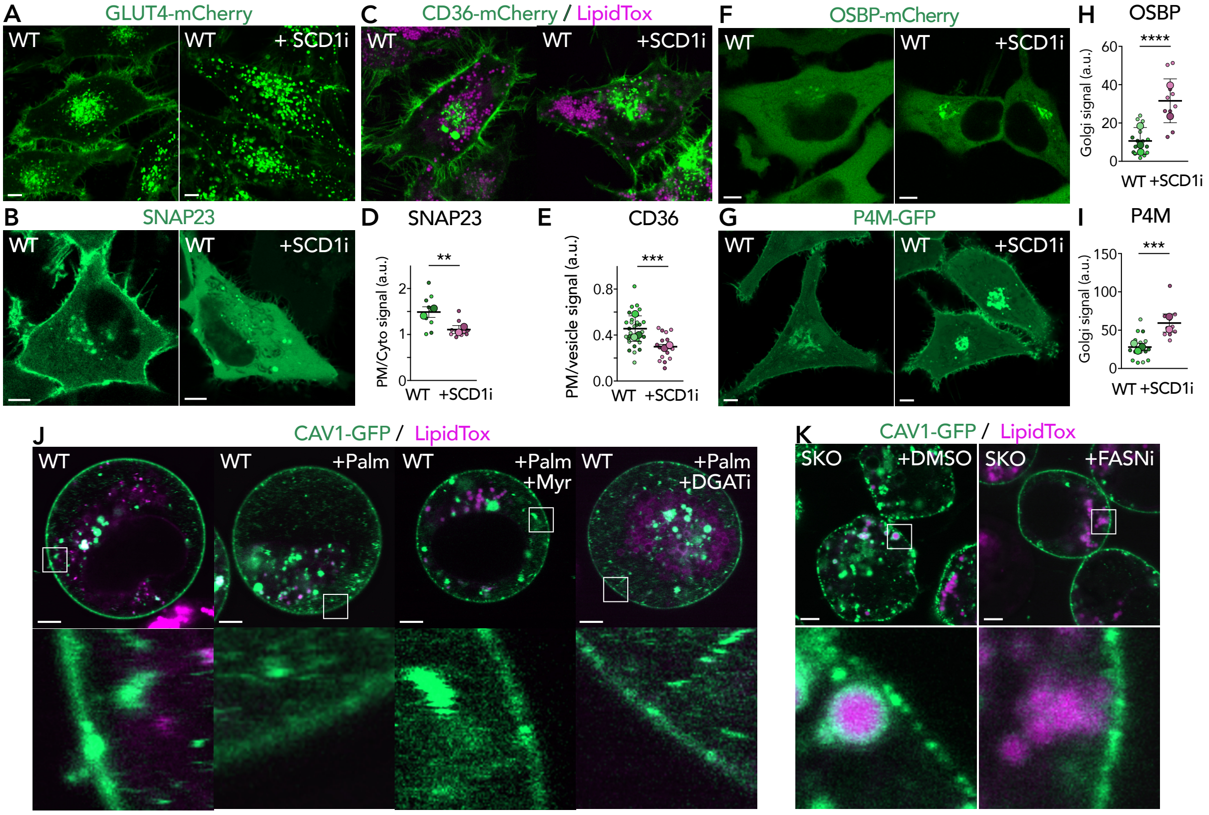


**Figure S8: A, B, C**. HeLa WT overexpressing GLUT4-mCherry, p-eGFPSNAP23, or CD36-mCherry. 1 µM of SCD1i is added 5 hours before transfection and maintained afterward. LDs are induced in CD36-mCherry transfected cells. **D, E.** Normalized PM mean signal for experiments in B and C. Panel D: Two-tailed Mann-Whitney analysis, P = 0.0089. n = 10 for WT and WT +SCD1i cells from 2 replicates. Panel E: P = 0.0006. n =30 for WT and 15 for WT + SCD1i from 3 and 2 replicates. **F, G.** HeLa WT overexpressing OSBP-mCherry or P4M-GFP. **H, I.** Quantification of experiments in F and G. Panel H: P < 0.0001. n = 15 for WT and 10 for WT + SCD1i cells, from 3 and 2 replicates. Panel I: Two-tailed Mann-Whitney analysis, P = 0.0004. n = 15 for WT and 10 for WT + SCD1i cells from 3 and 2 replicates. **J**. Swollen HeLa WT cells overexpressing CAV1-GFP, incubated with 2.5 µM of Myriocin (Myr) or 2.5 µM DGAT1/2 inhibitors (DGATi). After transfection, a palmitate/oleate mix is added for 24 hours. **K**. Swollen SKO LD-induced cells overexpressing CAV1-GFP. 1 µM of a FASN inhibitor (FASNi) is added 5 hours before the transfection and maintained afterward. Control is with DMSO. Scale bars: 5 µm.
